# Assessing the Functionality of RNA Interference (RNAi) in the Phloem-feeding Maize pest *Dalbulus maidis*

**DOI:** 10.1101/2021.09.29.462424

**Authors:** Tara-Kay L. Jones, Julio S. Bernal, Raul F. Medina

## Abstract

*Dalbulus maidis* [(DeLong & Wolcott), corn leafhopper], a phloem-feeding insect, is the most efficient vector of maize stunting pathogens (*Spiroplasma kunkelii*, Maize bushy stunt phytoplasma, and Maize rayado fino virus) in the Americas. Studies involving gene editing in insects are rapidly providing information that can potentially be used for insect vector and plant disease control. RNA interference (RNAi), a sequence-specific gene silencing method, is one of the most widely used molecular tools in functional genomics studies. RNAi uses exogenous double-stranded RNA (dsRNA) or small interfering RNA (siRNA) to prevent the production of proteins by inhibiting the expression of their corresponding messenger RNA (mRNA). In this study, we measured the efficacy of gene silencing, and its effects on *D. maidis* mortality as proof of concept that RNAi is a viable tool for use in genetic pest control of phloem-feeding insects. Oral delivery of dsRNA using an artificial diet was used to silence two key insect genes, vacuolar ATP synthase subunit B, and subunit D (*V-ATPase B* and *V-ATPase D*). Our results showed reduced gene expression of *V-ATPase B* and *V-ATPase D* after ingestion of dsRNA, and significantly higher mortality, and wing deformation, associated with reduced gene expression, compared to control insects that were not orally fed dsRNA. These results reveal RNAi as a viable tool for use in genetic pest control of phloem-feeding insects, and a way for further functional genomic studies, such as identification of potential target genes for either population suppression or population replacement of this vector of maize diseases.

## Introduction

*Dalbulus maidis* (DeLong & Wolcott) (Hemiptera: Cicadellidae), is a phloem-feeding, economically important pest of maize (*Zea mays* L. *mays*) (Coll-Aráoz et al., 2020; Meneses et al., 2016; Moya-Raygoza et al., 2014). *Dalbulus maidis* can cause up to 98% yield loss in maize by direct feeding on phloem sap and pathogen transmission (Nault & Bradfute, 1979; Pérez-López et al., 2018; Summers et al., 2004; Tsai, 2008). At least seven vector-borne diseases are known in maize, five of which are viral and two bacterial (Carloni et al., 2013; Hammond & Bedendo, 2005; Liu & Wang, 2018; Meyer & Pataky, 2010; Orlovskis et al., 2017; Zhang et al., 2011). *Dalbulus maidis* is the insect vector responsible for transmitting three of the most important maize disease agents, namely Maize rayado fino virus (MRFV), Maize bushy stunt phytoplasma, and *Spiroplasma kunkelii* (CSS), which it can acquire and transmit together or individually (Galvão et al., 2020; Mlotshwa et al., 2020; Sabato et al., 2020). Pathogens vectored by *D. maidis* are transmitted in a persistent, propagative manner, and as such *D. maidis* remains a vector throughout its life cycle, which facilitates disease spread across maize fields (Canale et al., 2018; Luft Albarracin et al., 2009; Ramos et al., 2020).

Currently, maize disease management relies on vector management with chemical pesticides, which poses health and environment risks (Nicolopoulou-Stamati et al., 2016; Özkara et al., 2016; Sabarwal et al., 2018; Singh et al., 2017). Alternatively, resistant maize germplasm against both *D. maidis* and the pathogens it vectors are available, but thus far are ineffective (Coll-Aráoz et al., 2020). Successful management of insect vectors and the pathogens they transmit may be achieved through genetic pest control. Genetic pest control is a strategy that utilizes molecular tools to manipulate pest genes, with the aim of reducing their population densities, or engineer refractory vectors for population replacement (Alphey & Bonsall, 2018; Scott et al., 2018). RNA interference (hereafter RNAi) is a promising tool with potential uses in both means of genetic pest control (Cagliari et al., 2019; Liu et al., 2020; Mamta & Rajam, 2017). RNAi is a sequence-specific method of silencing specifically targeted genes, which has already shown potential for disrupting pathogen transmission by vectors (Fletcher et al., 2020; Khare et al., 2018; Mahto et al., 2020; Pan et al., 2016; Petrick et al., 2016; Sammons et al., 2011; Younis et al., 2014).

RNAi was first discovered in the nematode *Caenorhabditis elegans* (Maupas) (Fire et al., 1998), and shortly thereafter its potential to control several insect pests was realized (Baum et al., 2007; Bramlett et al., 2020; Khajuria et al., 2018; Liu et al., 2019). This gene-silencing tool uses a mechanism where exogenous double-stranded RNA (dsRNA) inhibits the protein expression of targeted genes through degradation of corresponding messenger RNA (Mello & Conte Jr, 2004; Pecot et al., 2011). The basic principle for RNAi to be considered for pest control requires the delivery of dsRNA to insects through various means, including spraying of formulated concentrations directly on insects, transgenic plants capable of producing dsRNA, or nanoparticle delivery technologies (Andrade & Hunter, 2017; Baum et al., 2007; Christiaens et al., 2018; Ghosh et al., 2017; Jacques et al., 2020; Werner et al., 2020; Yan et al., 2020; Zha et al., 2011). Since 2008, private industry is exploring the development and application of RNAi for crop protection against insect pests (Bramlett et al., 2019; Head et al., 2017; Mat Jalaluddin et al., 2019).

Thus far, there are no published studies showing success of RNAi in *D. maidis*. However, oral delivery of dsRNA in other hemipteran (sap-sucking) insects, such as the stinkbug *Plautia stali*, was shown to be ineffective (Nishide et al., 2021). Effective delivery of dsRNA into insects is critical for widescale and commercial feasibility of RNAi based pest control. The functionality of RNAi has been shown in the corn planthopper, *Peregrinus maidis*, both through injection and oral dsRNA delivery methods, which silenced the vacuolar ATP synthase (*V-ATPase*) subunits (Yao et al., 2013). Likewise, the juvenile hormone acid O-methyltransferase (*HJAMT*), and vitellogenin (*Vg*) were successfully silenced by RNAi in the hemipteran brown marmorated stink bug (BMSB), *Halyomorpha halys*, through oral delivery of dsRNA via foliar sprays, root drenches, trunk injections, and clay granules (Ghosh et al., 2018). Similarly, in the hemipteran Southern green stinkbug, *Nezara viridula* L., a significant mortality of 45% was observed after orally inducing dsRNA gene silencing of *V-ATPase A* (Sharma et al., 2020). Finally, Galdeano et al. (2017) succeeded in inhibiting the expression of the *cathepsin D, chitin synthase*, an inhibitor of apoptosis genes, through oral delivery of dsRNA in the hemipteran Asian citrus psyllid, *Diaphorina citri* Kuwayama.

Several studies have assessed the functionality of RNAi in insect species by silencing gene subunits of the H^+^-translocating Vacuolar ATPase (V-ATPase) pump (Badillo-Vargas et al., 2015; Basnet & Kamble, 2018; Cooper et al., 2019; Thakur et al., 2014; Yao et al., 2013). The V-ATPase pump is an essential holo-enzyme for hydrolysis of ATP, a driver for nutrient uptake, and movement of protons out of cells to maintain membrane ion balance and cellular acidification of organelles, such as endosomes, lysosomes, and secretory vessels within insect organs. V-ATPase is found within the endomembrane and plasma membranes of insect cells, and is prevalent in the midgut, Malpighian tubules, and salivary glands. The V-ATPase synthase subunits play crucial roles in metabolic regulation in insects, and thus are widely used in assessments of RNAi gene silencing. For example, RNAi of V-ATPase B delivered by injection resulted in higher mortality and lower fecundity in the western flower thrips, *Frankliniella occidentalis*, most likely as a consequence of transcript and protein reduction (Badillo-Vargas et al., 2015). Given the conserved role of V-ATPases, the V-ATPase subunits B and D are ideal candidates for RNAi proof of concept, gene silencing in *D. maidis* through oral delivery using artificial diet.

Our main objective was to inhibit protein expression as proof-of-concept of the applicability of RNAi in *D. maidis*. dsRNA was delivered orally via an artificial diet to silence two key insect genes, vacuolar ATP synthase subunit B (*V-ATPase* B), and subunit D (*V-ATPase* D). Specifically, we measured the efficacy of gene silencing and its effects on *D. maidis* mortality and development to demonstrate that RNAi is a viable alternative for genetic pest control of this insect, as warranted. Proof of a functional RNAi machinery that is inducible by ingestion of dsRNA may accelerate the study of functional genes in *D. maidis* and related insects. Furthermore, a reliable method for orally delivering dsRNA into insect systems will facilitate research on molecular mechanisms of disease transmission and establishing that RNAi can be delivered orally can contribute to developing methods for managing insect vectors of plant diseases.

## Materials and Methods

### Insect rearing

*Dalbulus maidis* used in this study were maintained in colonies at the Texas A&M AgriLife Research and Extension Center in Weslaco, TX. The colonies were maintained on seedlings of a conventional maize hybrid, Anzu Genetica AG 1911, in their V4-6 growth stage in mesh cages, with a photoperiod of 14:10 (light: dark), and a temperature of 23-25 °C. The insects were originally collected in an experimental maize plot at 26.159244°, −97.960701°, and a commercial maize field at 26.083097°, −98.021841°, both in Weslaco, TX. Molecular identification of these leafhoppers was performed using the cytochrome oxidase I (COI) gene from five different individual insects; the corresponding sequences were deposited in the National Center for Biotechnology Information (NCBI) Genbank, under accession numbers MH591421 – MH591425.

### Target Genes-Primer design

RNAi efficiencies are well documented for evolutionarily conserved, and highly expressive target genes, such as vacuolar ATPases (V-ATPases) (Badillo-Vargas et al., 2015; Yao et al., 2013). *V-ATPase B* and *V-ATPase D* were used as target genes given their wide referencing and efficiencies as targets for RNAi. Nucleotide sequences of *Peregrinus maidis V-ATPase B* and *V-ATPase D* were extracted from GenBank in NCBI using accession numbers KC473163.1 and KC473162.1, respectively. These sequences were then blasted against a *D. maidis* transcriptome, generated and kindly provided by Dr. Astri Wayadande at Oklahoma State University, to identify the homologs in *D. maidis* (Wayadande et al., unpublished). The E-RNAi tool (https://www.dkfz.de/signaling/e-rnai3/idseq.php) was used to generate two primer pairs for each gene of interest, as well as sequences of the green florescence protein ([pBIN-mGFP]-for experimental controls). Forward and a reverse primer containing the T7 promotor sequence TAATACGACTCACTATAGGG, and additional forward and reverse primers without the T7 promotor sequence were generated (Table 1). Furthermore, real-time qPCR primers to validate the effectiveness of *V-ATPase-B* and *V-ATPase* D silencing were generated (Table 2.) using the Primer3 tool (http://bioinfo.ut.ee/primer3-0.4.0/), with parameters adjusted to exclude the dsRNA targeted regions of both *V-ATPase* genes of interests. The efficiency of each of the primer sets was determined by means of qRT-PCR using *D. maidis* cDNA dilutions of 2-folds.

**Table 1.**
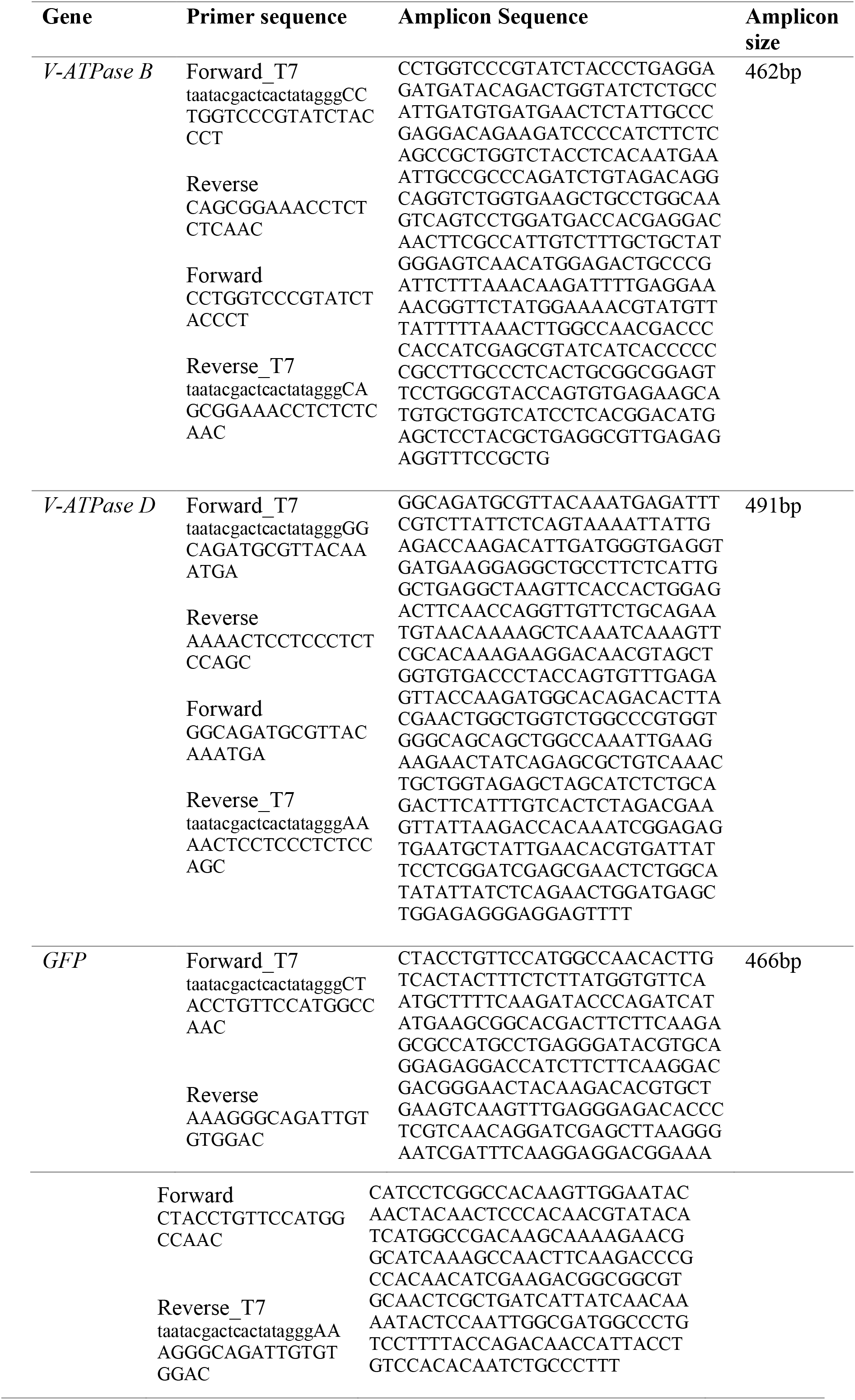
Sequences for dsRNA synthesis of V-ATPase B, V-ATPase D and GFP

**Table 2.**
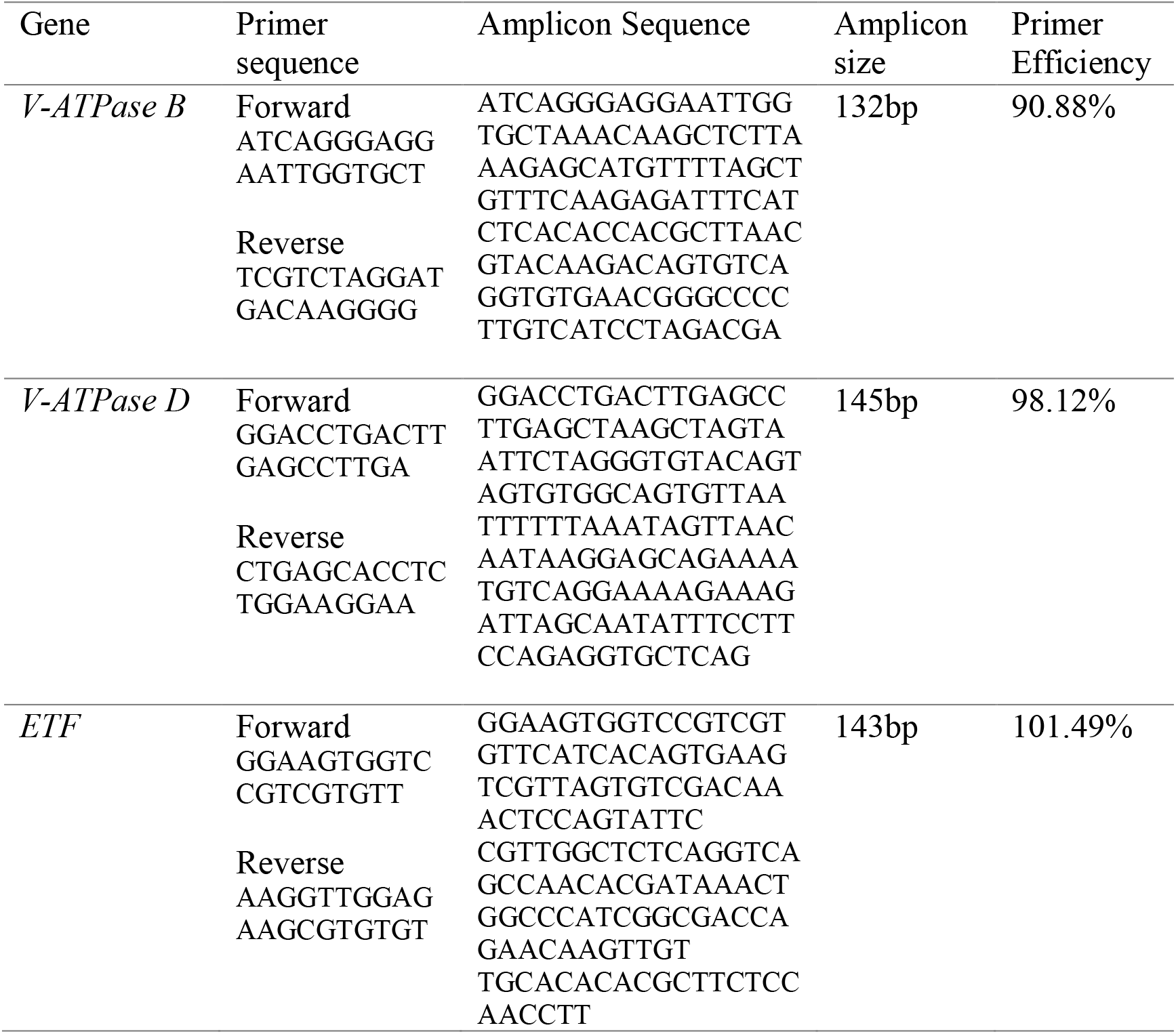
Sequences for qRT-PCR evaluation of V-ATPases silencing in *D. maidis*.

### dsRNA Synthesis

Total RNA was extracted from a pool of twenty (20) *D. maidis* adults using a TRIzol extraction method (Badillo Vargas et al., 2012). One microgram of total RNA was used to synthesize cDNA using the Verso cDNA Synthesis Kit (Thermo Scientific^™^, MA, USA) according to the manufacturer’s protocol. Polymerase chain reaction (PCR) was performed to amplify gene products from *D. maidis V-ATPase* B and *V-ATPase* D cDNA. For maximization of gene product yield, two separate PCR reactions for each gene target and control were carried out (*i.e*., primer pair Fwd_T7 + Rev were combined in one reaction followed by a separate reaction with Rev_T7 + Fwd). The GFP dsRNA was amplified from a plasmid DNA. PCR reactions were run with a hot start of 94 °C for 5 min, followed by 9 cycles of denaturation at 94 °C for 30 sec, annealing of gene specific primers at 55 °C for 30 sec, and extension at 72 °C for 1 min, followed by a second 34 cycles of denaturation at 94 °C, annealing of T7 primers at 65 °C, and an extension at 1 °C in a Bio-RAD T100^™^ Thermal Cycler (Bio-RAD, Hercules, CA). PCR amplicons from each reaction were purified using a QIAquick PCR Purification kit (Qiagen, Venlo, Netherlands). *In vitro* transcription to generate dsRNA for each target gene and the negative control was performed by combining reagents of the T7 RiboMAX^™^ Express RNAi System (Promega, Fitchburg, WI) according to the manufacturer’s instructions. Briefly, reactions were incubated at 37 °C for 30 min. Both products per specific gene were combined to form the dsRNA at 70 °C for 10 min, then cooled at room temperature for 20 min. dsRNA was then treated with single units of RNase solution, and of RNA Qualified DNase (RQ1), followed by incubation at 37 °C for 30 min. Each dsRNA target was further purified using 3M sodium acetate, and one volume of isopropanol, incubated on ice for 5 min, and centrifuged at high speed (12,000 rpm) for 10 min. The supernatant was discarded, and the dsRNA pellets were washed twice with 70% ethanol, and air dried at room temperature for 15-30 min, after which dsRNA was resuspended in 20 μL of nuclease-free water, followed by quantification using a Nanodrop, and stored at −80 °C until further use.

### RNAi knockdown of V-ATPase B and D through feeding

A time-course experiment of 4 days was conducted to determine the effect of oral delivery of dsRNA on expression of relevant genes, survivorship after 4 days, and wing development in *D. maidis*. The experiment included four treatment groups: (i) dsRNA V-ATPase B, (ii) dsRNA V-ATPase D, (iii) dsRNA GFP (= control for dsRNA consequences because GFP is absent in insects), and (iv) buffer (= control lacking dsRNA, containing only nutrients). Each treatment had three experimental replicates, randomly arranged, and kept in insect mesh cages at 14:10 (light: dark) photoperiod and a temperature of 23-25 °C. Treatment groups comprised third instar *D. maidis* nymphs held within feeding chambers constructed by modifying 50ml graduated polypropylene centrifuge tubes (Thermo Fisher Scientific Inc. USA) to be 3cm (diameter) × 5.5cm (length), with insect-proof nylon mesh on the underside, and sealed by parafilm to prevent insects from escaping. Preliminary assays showed that third instar nymphs showed the highest and most uniform expression level of both *V-ATPase B* and *D*, so selected as targeted stage for RNAi (unpublished data). A final volume of 100μl of D-10 feeding solution (containing 10% Sucrose, 0.2% Fructose, 0.38% Potassium Phosphate, 0.03% Magnesium Chloride and 1% Fetal bovine serum, pH=7) (Carpane et al., 2011), and 0.5μg/μL (500 ng/μL) dsRNA was placed between two layers of parafilm, which was the ceiling of the feeding chambers. Each treatment diet was replenished every 24 h to maintain stability and integrity of the dsRNAs in the feeding solution. Three alive insects were collected at 2- and 4-days post-feeding from each treatment groups (*i.e*., n = 3 independent replicates, per treatment per time point [2- and 4-days], for a total of 24 samples). The insects were stored at −80 °C, and later processed for total RNA and qRT-PCR of gene silencing.

### Quantification of gene expression

As noted above, groups of three alive insects were collected at 2- and 4-days post feeding on 2 dsRNA treatments (i.e., dsRNA V-ATPase-B and D) and 2 control groups (i.e., dsRNA GFP and buffer diet only) for total of 24 pooled samples (each pool consisted of 3 insects, thus a total of 72 insects used for the analysis) and processed for real-time quantitative reverse-transcriptase PCR (qRT-PCR) analysis. Total RNA was extracted from each pool of insects collected from the dsRNA assay described previously in RNAi knockdown of *V-ATPase B* and *D* through feeding section, using Trizol reagents (Badillo Vargas et al., 2012). RNA quantity was analyzed by Nanodrop Spectrophotometer (Thermo Fisher Scientific Inc. USA) at 260 nm. The first strand cDNA of each pool was synthesized from 1μg total RNA using the ThermoScientific Verso cDNA synthesis kit with RT-enhancer (ThermoScientific Inc. Waltham, MA) to remove residual DNA contamination. A single qRT-PCR reaction consisted of 4 μL of cDNA (diluted 1:10), 300nM final volume of each forward and reverse primer pair, 5 μL SYBR Green Supermix (50% total volume of reaction, rxn) (Bio-Rad Inc. Hercules, CA), and nuclease-free water (to make a final reaction volume of 10 μL), +10% extra volume to allow for pipetting error. The qRT-PCR was performed in a 2-step amplification with 40 cycles of 95 °C for 30 s and 55 °C for 30s using a Bio-Rad IQ thermocycler (Bio-Rad Laboratories Inc. Hercules, CA). Each reaction was loaded on a 384-wells standard qPCR plate and ran in triplicates. The normalized abundance of target transcripts (*V-ATPase B* and *D*) to the internal reference transcript (*ETF*) was calculated for the treatment (dsRNA V-ATPases) and the control (dsRNA GFP) samples. The relative abundance of target transcripts (*V-ATPase B* and *D*) in the treatment (dsRNA V-ATPase) compared to the control (dsRNA-GFP) was calculated using the 2–ΔΔCT algorithm (also known as the delta-delta-Ct or ddCt algorithm) (Livak & Schmittgen, 2001).

### Effect of oral delivery of dsRNA V-ATPase B and D on nymphs’ survival

A separate time-course experiment with four treatment groups, dsRNA V-ATPase B, dsRNA V-ATPase D, dsRNA GFP control, and a buffer control, were conducted over a 4-day period to determine the effect of oral delivery of dsRNA on survival of *D. maidis*, relative to the controls (dsRNA GFP, buffer). Groups of 30 fifth instar *D. maidis* nymphs were held in each of two feeding chambers, each of which represented one replicate of each diet. All diet treatments were replenished daily to maintain treatment integrity and avoid starvation (Yao et al., 2013). The experiment was repeated three times, for six total replicates; overall, 720 insects were used to evaluate the effects of oral delivery of dsRNA on *D. maidis* survival and wing development. The numbers of living insects and insects with deformed wings were counted after 4 days for each treatment.

### Statistical Analysis

The effects of dsRNA oral delivery on V-ATPase B or V-ATPase D expression in *D. maidis* were assessed by analyses of variance (ANOVA) of ratios of V-ATPase subunit expression to dsRNA GFP expression (= expression in V-ATPase subunit treatment/expression in GFP control); ratios were log10-transformed for normality prior to analysis (Shapiro-Wilk *W* test stats, P≥ 0.202). The ANOVA model included the independent variables V-ATPase subunit (B or D) and day (2 and 4), and the interaction term (V-ATPase subunit * day); *a priori* contrasts were used to separate per-day means within V-ATPase subunits, as warranted. One-sample *t* tests were used to test the hypothesis that the mean ratio for each of the V-ATPase subunit * day pair did not differ from 1.0 (i.e., expression in V-ATPase subunit treatment/expression in GFP control = 1.0); a critical P value of 0.013 was used for each *t* test, per Bonferroni’s correction (Abdi, 2007). The effects of dsRNA treatments and control on *D. maidis* survival after 4 days were assessed using ANOVA, followed by Tukey’s tests to compare dsRNA control means among V-ATPase subunits; survival rates (= insects alive at day 4/insects alive at day 0) were arc-sine square-root √*x*-transformed prior to ANOVA. Finally, the effects of dsRNA treatments on wing development were assessed by comparing frequencies of individuals with deformed wings using Chi-square tests with Yates’ correction for continuity (Upton, 2000). All statistical analyses were conducted using JMP PRO 15.0.0 (SAS Institute Inc, NC.USA).

## Results

### Effect of oral delivery of dsRNA on V-ATPase-B and D expression in third-instar nymphs

Insects fed with diets containing dsRNA V-ATPase B and D exhibited a reduction in normalized transcript abundance compared with controls that fed on diets containing dsRNA GFP (a non-target gene sequence absent in insect). ANOVA did not reveal gene (V-ATPase B or D) (P = 0.583), day (2 or 4) (P=0.403), or gene*day interaction (P=0.680) effects on mRNA transcript levels (Fig. 1). However, one-sample *t* tests showed that mRNA transcript levels were significantly reduced below baseline level (ratio < 1.0) on day 4 in both *D. maidis* fed diet containing dsRNA V-ATPase B or D (P= 0.012 and 0.01, respectively) (Fig. 1). These results indicated that dsRNA V-ATPase B and D triggered RNAi gene silencing in *D. maidis* after 4 days of ingestion.

**Figure 1.**
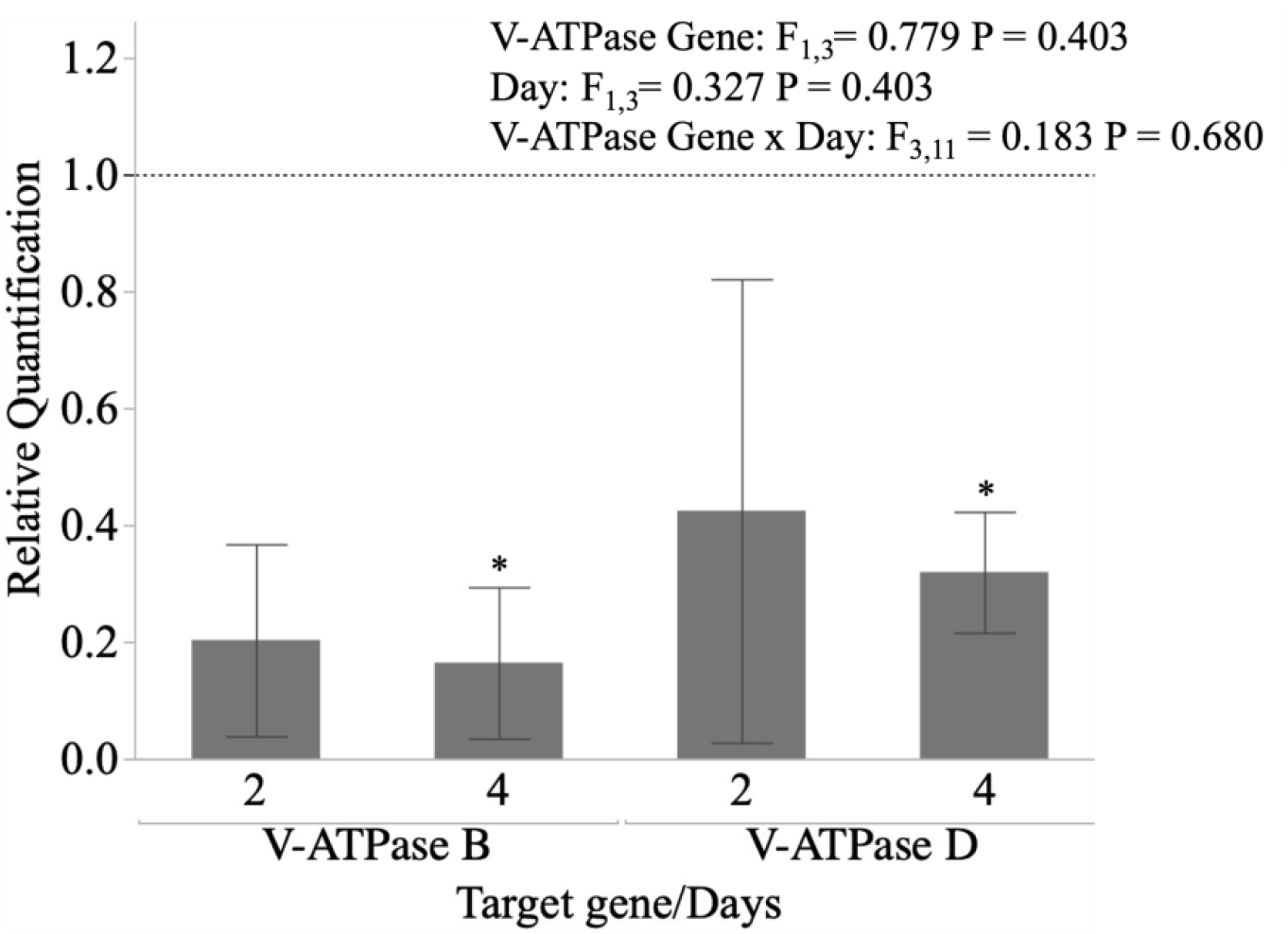
Effect of RNAi knockdown on gene expression in third-instar nymphs. Expression levels (shown as ratios) of *V-ATPase B* and *D* genes in *Dalbulus maidis* at 2 and 4 days post oral delivery. Expression levels are shown relative to expression of GFP control; thus, ratio value 1.0 indicates no difference in expression levels between V-ATPase and GFP control. ANOVA stats are shown inset; asterisks above columns indicate significant difference from ratio 1.0 for V-ATPase B day 4 (df= 2, P ≤ 0.012) and V-ATPase D day 4 (df= 2, P≤ 0.011).

### Effect of oral delivery of dsRNA V-ATPase B and D on nymphs’ survival

Survival of *D. maidis* was lowest in the *V-ATPase B* and *D* treatments, and highest in the GFP and no dsRNA controls (F= 20.763; df=3, 23; P < 0.0001) (Figure 2). Survivorship began decreasing at ca. 2 days in the no dsRNA treatment, and prior to 2 days in the remaining treatments. At 4 days, survival was ca. 52% in the no dsRNA treatment, and ~37%, ~18%, and ~16% in the GFP, V-ATPase D, and V-ATPase B treatments, respectively. The similarity between survival rates in the no dsRNA and GFP control treatments in contrast with the similar and significantly lower survival rates in the V-ATPase D and B treatments suggested that the decreased survival in the latter two treatments was due to RNAi knockdown.

**Figure 2.**
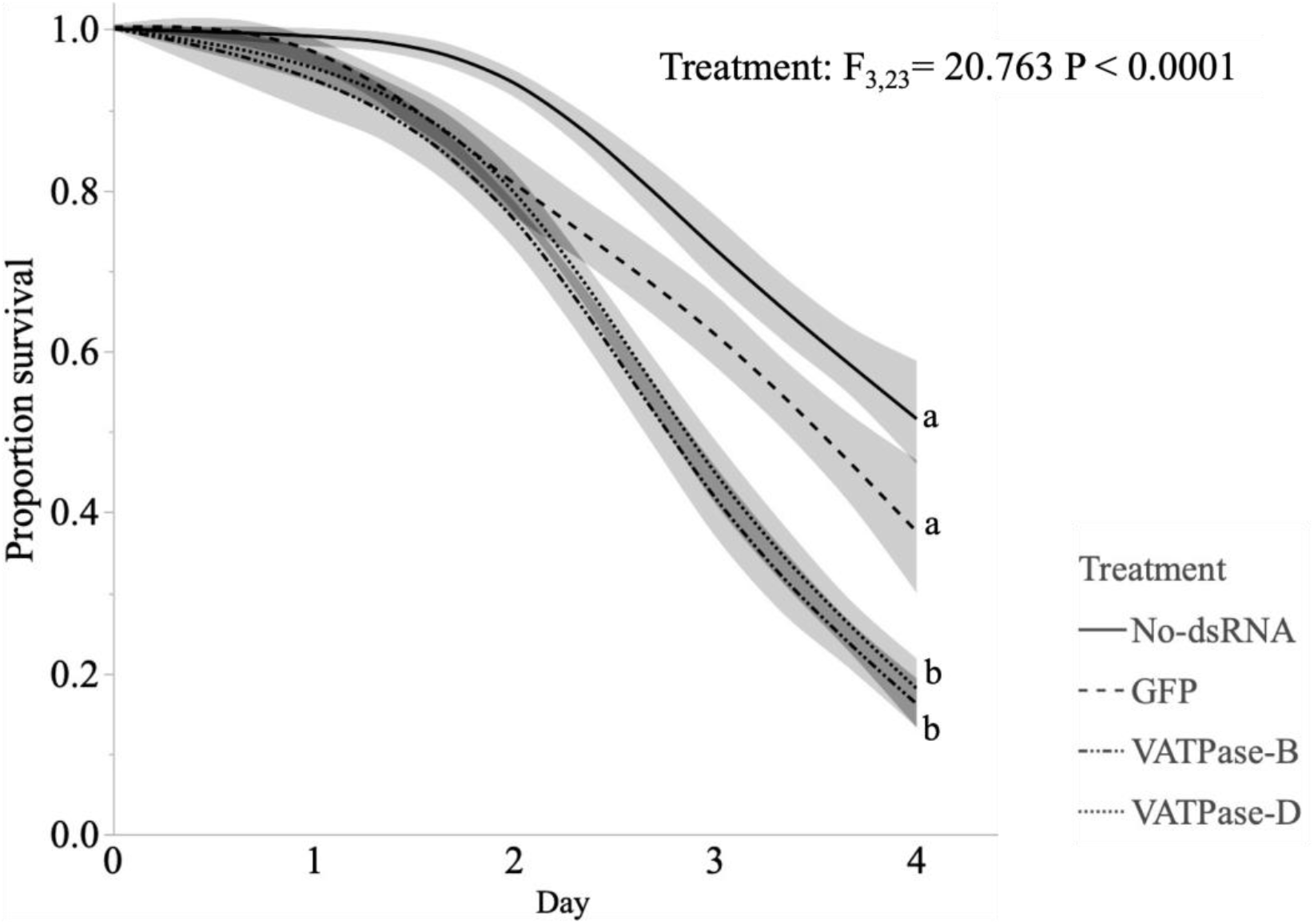
Effect of RNAi knockdown on fifth-instar nymph’s survival. Survival (shown as proportions) of *Dalbulus maidis* fifth instar nymphs after 4 days post oral delivery of dsRNAs. ANOVA stats are shown inset; connecting lower-case letters (a, b) represent statistical differences among LSMeans of treatments (levels not connected by the same letter represent statistical significance) by Tukey HSD (α= 0.050, Q= 2.799). Smoothers around fit line of each treatment represents confidence of fit.

### Effect of oral delivery of dsRNA V-ATPase B and D on emerging adult’s phenotype

*Dalbulus maidis* adults that molted from fifth instar nymphs during dsRNA V-ATPase B and D treatment typically developed with deformed and small wings showing excessive curling and resulting in reduced mobility (Fig. 3a). A Chi-square test revealed statistical significance in the frequency of deformed adults that molted after dsRNA ingestion of V-ATPase B (χ^2^ critical =6.897; P= 0.008). No statistical significance was detected in the frequency in deformed occurrences due to *V-ATPase D* knockdown by chi-square (χ^2^ critical =2.130; p-Value =0.144) (Figure 3b). The hind wings were notably more affected than the forewings, consistent with the severe reduction in mobility and flight.

**Figure 3.**
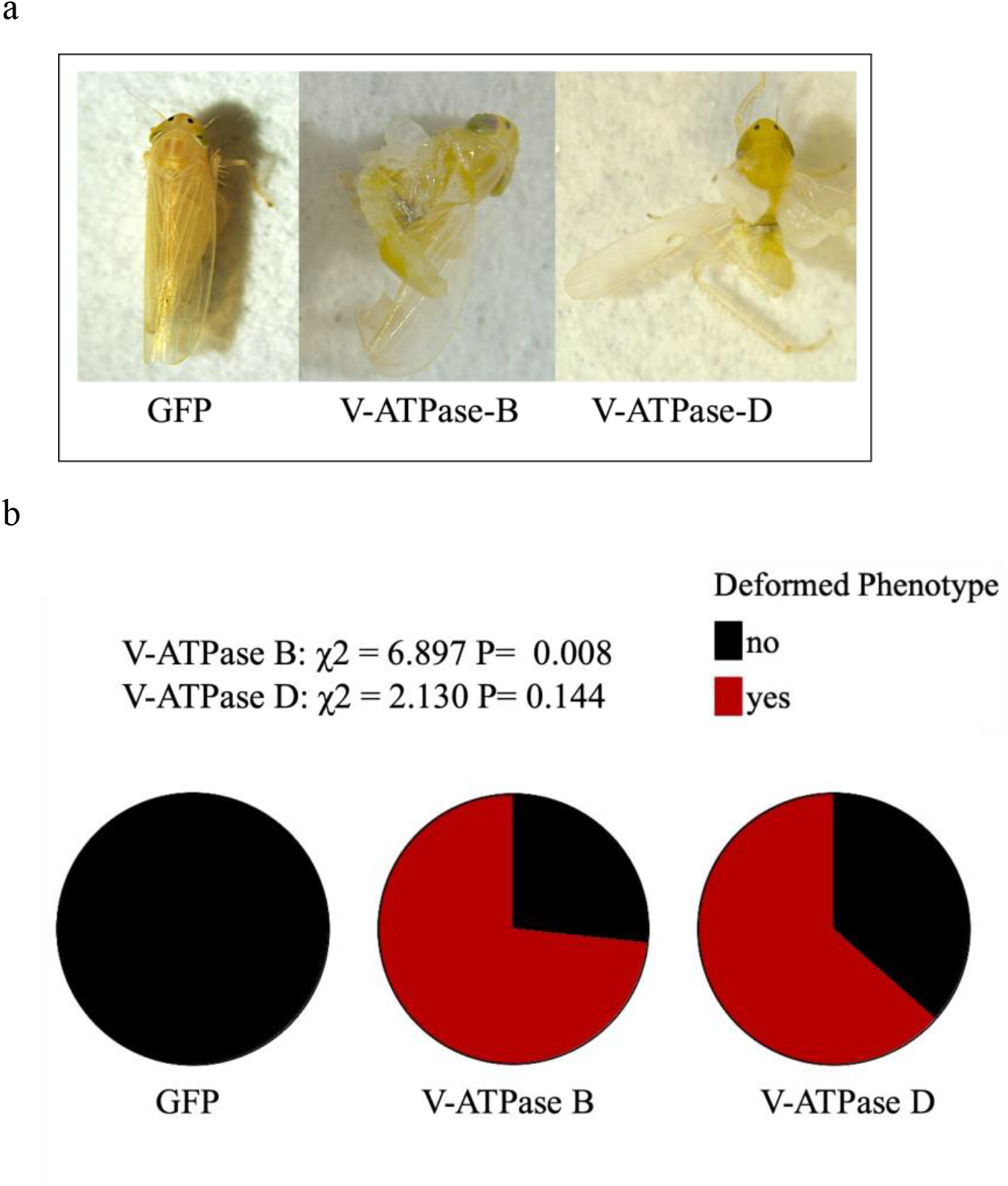
Effect of RNAi knockdown on emerging adult’s phenotype. (a) Deformed wings and abdomen of *Dalbulus maidis* upon molting 4 days post oral delivery of dsRNAs. (b) Proportion of deformed insect after fifth instar nymphs molted into adults. Chi-square stats are shown inset.

## Discussion

Our results showed that oral delivery of dsRNA causes partial silencing of *V-ATPase B* and *D* genes and is associated with higher mortality and greater frequency of wing deformations in *D. maidis*. Thus, our results suggest that RNAi may be a viable alternative for management of hemipteran (phloem-feeding) pests. Oral delivery of dsRNA led to decreased expression of *V-ATPase B* and *D* genes in *D. maidis*, relative to their expression level when compared to effects of green florescence protein (pBIN-mGFP) dsRNA, as shown by qRT-PCR assays, which was correlated with increases in mortality and wing malformations during molting. Also, we observed that silencing was initiated by 2 days post treatment and was significant after 4 days.

Structurally, the V-ATPase is a rotary nanomotor which comprises of two domains: a peripheral V1 domain (Subunits A-H) and a membrane intrinsic V0 domain (subunits a, c, d, and e) (McGuire et al., 2017; Wieczorek et al., 2009). The gene targets selected for silencing in our study, V-ATPase subunits B and D, both make up the V1 domain, which plays a role in ATP hydrolysis, a process necessary for releasing energy used in cellular processes. Synthesis of all subunits are required for functionality of the V-ATPase pump, so disruption of a single unit affects its overall assembly. Therefore, the significant reduction in gene expression of both *V-ATPase B* and *D* are positive indication that the insects were likely experiencing disruption of normal hydrolysis of ATP, which could cause membrane ionic imbalances, disruption in nutrients absorption, and metabolic deregulation. The success in RNAi knockdown of V-ATPase subunits causes inhibition of energy release through ATP, thus impairing necessary biological functions, and likely leading to the >50% mortality we recorded in *D. maidis* at 4 days post ingestion of dsRNA.

RNAi knockdown of V-ATPase B crosslinks with several other V-ATPase subunits, such as E, G, and C to form the peripheral stalk of the proton pump (Basnet & Kamble, 2018; Nishi & Forgac, 2002). The peripheral stalk is also comprised of V-ATPase subunit D, thus, silencing of V-ATPase B may have induced an inhibitory expression of subunit D, and all other subunits, resulting in non-functionality of the entire proton pump. The disruption of functionality of such important enzyme (V-ATPase) may explain the wing deformity frequently associated with dsRNA ingestion in *D. maidis*. Numerous studies have been conducted on V-ATPase in insects and other arthropods with regards to molting and metamorphosis (Hou et al., 2020; Phillips, 2002; Rahmani & Bandani, 2021). In hemimetabolous insects, such as *D. maidis* and hemipterans, generally, molting is associated with wing growth. We observed frequent deformity in wings of *D. maidis* insects as they molted into adults from the fifth instar, nymphal stage. We suggest that wing development, and growth during the molting process, may have been hindered due to disruptions in the assembly of the V-ATPase pump. Studies have reported that in *Drosophila melanogaster* fruit flies, and *Manduca sexta* caterpillars V-ATPase interacts with protein kinase A, which phosphorylates protein substrates to regulate its assembly/disassembly and facilitate energy utilization during molting (McGuire et al., 2017; Voss et al., 2007). Molting is an energy intensive process (Arrese & Soulages, 2010; Lorenz & Gäde, 2009), so limitation of internal cellular energy from ATP can significantly compromise developmental processes, such as cuticular formation, and wing development (Hou et al., 2020). Many of the RNAi-treated insects that underwent molting in our experiment did not develop hardened exoskeletons in the thoracic region and showed abnormal sclerotization and pigmentation (data not shown). Silencing of V-ATPase subunits resulted in similar molting deformities in other insect species, such as *Periplaneta fuliginosa*, the smokey brown cockroach (Sato et al., 2019), *Phenacoccus Solenopsis*, the cotton mealybug (Khan et al., 2018), *Locusta migratoria manilensis*, the Oriental migratory locust (Li & Xia, 2012), and *Henosepilachna vigintioctopunctata*, the 28-spotted potato ladybird (Zeng et al., 2021), among others (Badillo-Vargas et al., 2015; Nishide et al., 2021; Yao et al., 2013).

No prior published studies have showed RNAi in *D. maidis*. However, the partial silencing that we found is consistent with previously published research on the related species *Peregrinus maidis*, corn planthopper. Similarly, RNAi knockdown of *V-ATPase B* and *D* was not significant in *P. maidis* at 2-days post ingestion, whereas the effects of gene silencing statistically significant in feeding after 4-days or more (Yao et al., 2013). In other hemipteran species such as *Cimex lectularius* L., the common bed bug, RNAi of V-ATPase subunits significantly reduced mRNA transcript as early as 3-days of treatment (Basnet & Kamble, 2018). As molecular methods improve, more studies are reporting successful oral delivery of dsRNA molecules overcoming the known hemipteran cellular barriers against RNAi (Ghosh et al., 2018; Jain et al., 2021; Sharma et al., 2021). Evidence of successful RNAi mechanism induced by oral delivery of dsRNA was recently presented in *Pseudococcus maritimus*, the grape mealybug (Arora et al., 2021). As in our study, RNAi silencing was evident in *P. maritimus* as early as 3-days when insects were fed dsRNA complements of osmoregulation genes (aquaporin (*AQP*) and a sucrase (*SUC)*) (Arora et al. 2020).

## Conclusion and Future Prospective

This study is the first to show as proof-of concept that RNAi is a functional mechanism in *D. maidis* and thus can serve as an ideal model system for comprehensive genome studies. Partial silencing of V-ATPase subunits B and D resulted in significant reductions in corresponding gene expression, higher mortality, as developmental malformations in *D. maidis*. RNAi is typically administered through microinjection of dsRNA into the insect body. The main advantage of microinjections is that it bypasses salivary and midgut barriers, including dsRNAs, which reduce sensitivity, and affect oral delivery methods (Allen & Walker Iii, 2012; Christiaens et al., 2014). However, RNAi by microinjection is not a practical method for pest control use. In our experiments we showed that dsRNA uptake was successful through an oral ingestion strategy using artificial diet, which suggests that in *D. maidis* dsRNA may bypass salivary and midgut barriers, and access cells to disrupt gene expression and confer systemic responses through reduced expression of targeted genes. Whilst artificial diets are not ideal delivery method for pest and disease vector control in real-word settings, our study shows that oral ingestion could be effective for RNAi uses. For example, could be delivered to plant feeding pests via in-planta systems, or generally via nano-pesticide formulations. Thus far, RNAi has been induced via nanoparticle formulations which administered dsRNA to agricultural insect pests, such as *Chilo suppressalis*, the striped rice stemborer (Zhang et al., 2021), *Aphis glycine*, the soybean aphid (Yan et al., 2020; Zhang et al., 2021; Zheng et al., 2019), and *Myzus persicae*, the green peach aphid (Zhang et al., 2021). The latter two pests, like *D. maidis*, are hemipterans.

*D. maidis* is the most important vector of the pathogens causing several important maize diseases. Controlling maize diseases relies on understanding the interactions between insect vectors and pathogens transmission. A successful RNAi strategy can also be useful in further dissecting the molecular interactions that mediate the role of *D. maidis* in pathogen transmission by facilitating the study of key genes and their functions within the insect. Insect proteins and enzymes play critical roles in the transmission of vector-borne plant pathogens. For example, *Spiroplasma citri*, the causative agent of citrus stubborn disease, glycolytic enzyme phosphoglycerate kinase (PGK) is known to bind *Circulifer haematoceps* actin for cellular internalization necessary for transmission (Labroussaa et al., 2010; Labroussaa et al., 2011). To date, there is a shortage of studies addressing the roles of vector derived proteins in *D. maidis* transmission of maize pathogens. Furthermore, the *D. maidis* pathosystem is an excellent choice for comprehensive studies of vector-borne plant diseases because it transmits both bacterial and viral pathogens (Jones & Medina, 2020). Our study paves the way for elucidating the roles of gene candidates in vector transmission. Upon identification of key gene targets in *D. maidis* pathosystem, RNAi can then be applied as a strategy in the disrupting interactions between pathogen and vector through gene suppression, which may lead to reduction of disease spread.

## Acknowledgement

We would like to thank the Texas AgriLife Research and Extension Center at Weslaco for supporting this research, and Ismael E. Badillo (formerly at Texas AgriLife Research and Extension Center at Weslaco) for editing the early versions of this manuscript.

